# Specific features and assembly of the plant mitochondrial complex I revealed by cryo-EM

**DOI:** 10.1101/2020.02.21.959148

**Authors:** Heddy Soufari, Camila Parrot, Lauriane Kuhn, Florent Waltz, Yaser Hashem

## Abstract

Mitochondria are the powerhouses of eukaryotic cells and the site of essential metabolic reactions. Their main purpose is to maintain the high ATP/ADP ratio that is required to fuel the countless biochemical reactions taking place in eukaryotic cells^1^. This high ATP/ADP ratio is maintained through oxidative phosphorylation (OXPHOS). Complex I or NADH:ubiquinone oxidoreductase is the main entry site for electrons into the mitochondrial respiratory chain and constitutes the largest of the respiratory complexes^2^. Its structure and composition varies across eukaryotes species. However, high resolution structures are available only for one group of eukaryotes, opisthokonts^3–6^. In plants, only biochemical studies were carried out, already hinting the peculiar composition of complex I in the green lineage. Here, we report several cryo-electron microscopy structures of the plant mitochondrial complex I at near-atomic resolution. We describe the structure and composition of the plant complex I including the plant-specific additional domain composed by carbonic anhydrase proteins. We show that the carbonic anhydrase is an heterotrimeric complex with only one conserved active site. This domain is crucial for the overall stability of complex I as well as a peculiar lipid complex composed cardiolipin and phosphatidylinositols. Moreover we also describe the structure of one of the plant-specific complex I assembly intermediate, lacking the whole P_D_ module, in presence of the maturation factor GLDH. GLDH prevents the binding of the plant specific P1 protein, responsible for the linkage of the P_P_ to the P_D_ module. Finally, as the carbonic anhydrase domain is likely to be associated with complex I from numerous other known eukaryotes, we propose that our structure unveils an ancestral-like organization of mitochondrial complex I.

---

Complex I is the largest multimeric enzyme of the respiratory chain, composed of more than 40 protein subunits. 14 are strictly conserved proteins subunits, vestige of complex I of bacterial origin^7^. The additional subunits, referred to as supernumerary subunits, were acquired during eukaryotes evolution. Part of these additional subunits are conserved among other eukaryotes and play essential roles for the structure, function and the association with the other respiratory chain complexes, eg. to assemble into respirasome in animals^8^. Recently, several high resolution 3D structures of the complete mitochondrial complex I of opisthokonts were determined by cryo-EM in mammalian species^4–6^ and in the aerobic yeast *Yarrowia lipolytica*^3,9^, revealing the organization of their additional specie-specific subunits. However, in plants, even though extensive biochemical characterization was conducted^10,11^, high-resolution structures of mitochondrial complex I are yet to be derived. Early negative staining studies^12^ revealed the presence of a large additional membrane attached domain, absent from animal and yeast species, hinting at the peculiar structure and composition of the plant complex I.

In order to obtain a high-resolution structure of the plant mitochondrial complex I and its additional subunits, we purified mitochondria from *Brassica oleracea var. botrytis*, a close relative to the model plant Arabidopsis (both belong to the group of Brassicaceae plants), as previously described^13^ (see Methods). We recorded cryo-EM images of membrane complexes purified from sucrose gradient (see Methods), corresponding to mitochondrial complex I. After particle sorting (see Methods) we obtained cryo-EM reconstructions of two main complex I states: the full plant mitochondrial complex I as well as reconstruction of a complex I assembly intermediate, without the PD module, and with the plant-specific assembly factor GLDH^14^ (Extended Data Fig. 1). Focused refinement and particle polishing were performed in RELION3^15^ (see Methods), allowing reconstruction of the full complex I at 3.7Å resolution, focused classification on the membrane arm and P_P_ module yielding 3.4Å. The P_D_ module being more flexible produced more scant densities reporting lower resolutions (Extended Data Fig. 1 and 2). The cryo-EM maps allowed the determination of many well-resolved features and clear side-chain densities (Extended Data Fig. 3) that enabled modelling of the 45 proteins of the plant mitochondrial complex I (Extended Data Table 1), as well as several characteristic ligands that were directly identified from the density: the eight canonical FeS clusters of the matrix arm, the FMN of the 51kDa subunit, the NADPH molecule in the 39kDa, three cardiolipins two phosphatidylinositol and one phosphatidylethanolamine (Extended Data Fig. 3). Ubiquinone was also clearly visible in the Q module (Extended Data Fig. 4).

## General description

The plant mitochondrial complex I has the classical open L-shape formed by the matrix and membrane arms (Fig. 1). In the matrix arm, electrons are transferred from NADH to ubiquinone along the FeS clusters with distances similar to what has been described in bacteria and opisthokonts mitochondrial complex I (Extended Data Fig. 6). The membrane part, where proton pumping takes place, is also conserved. Those highly conserved features are part of the 14 minimal protein subunits conserved in bacteria and other mitochondrial complex I^2^. The additional mitochondria specific proteins enhance the volume and mass of the complex I compared to bacteria. In plant, 25 proteins shared with aerobic yeast or mammals are present. Similarly to the previously obtained complex I structure^4,6,9,16^, the supernumerary subunits, specific to mitochondrial complex I, form a shell around the core subunits, adding nearly 400kDa of proteins to the conserved subunits (Extended Data Fig. 6). These supernumerary subunits are mainly composed of α-helices and are largely interconnected with the core subunits. In the Q module of the matrix arm, our reconstructions allowed us to clearly visualize ubiquinone in the pocket formed by Nad1, PSST and Nad7 (Extended Data Fig. 4). Our results confirm the recently observed ubiquinone position described for *Y.lipolytica*^3^.

**Figure 1.**
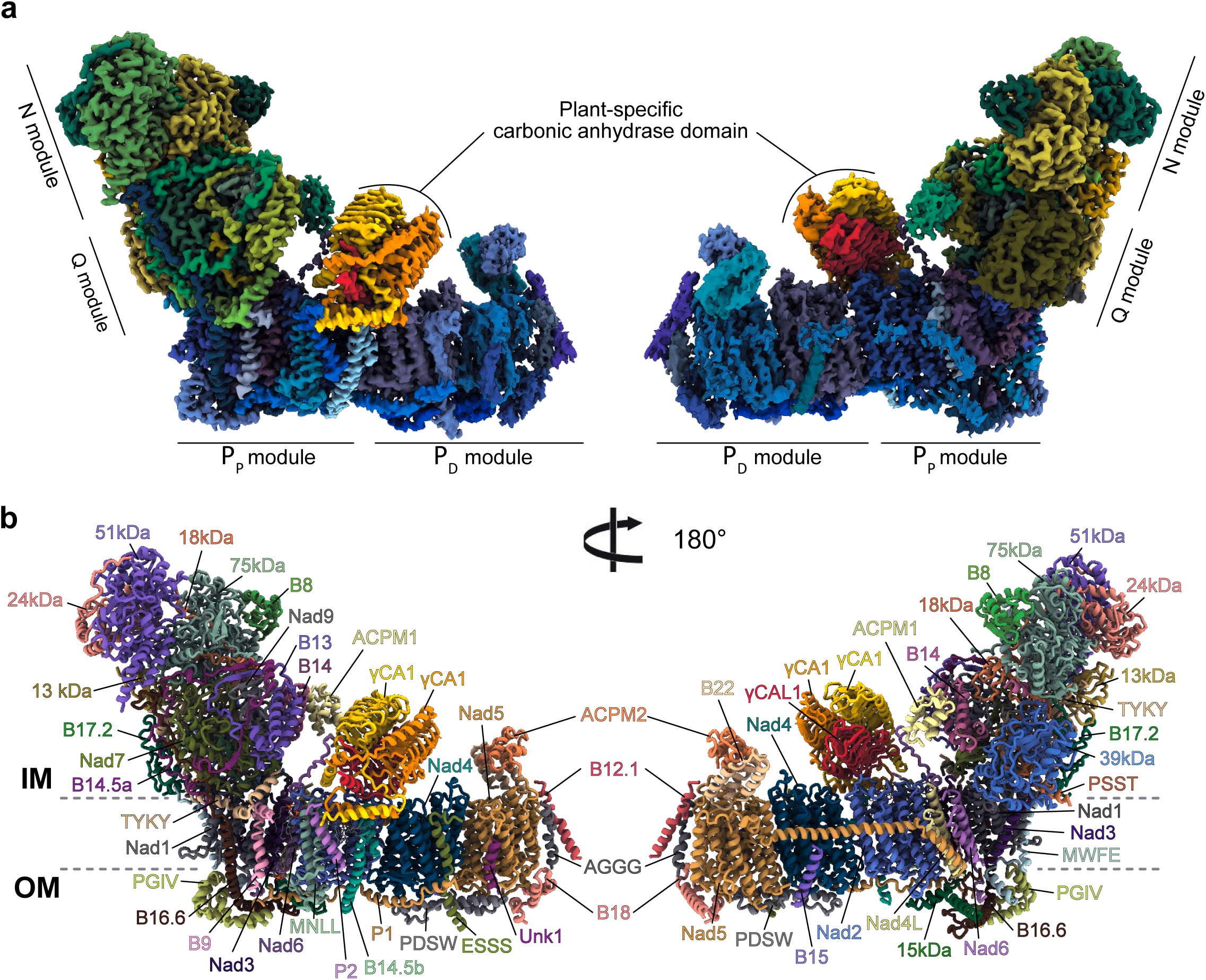
Overall structure of the plant mitochondrial complex I. **a** Composite cryo-EM map of the complete plant complex I. The membrane part is displayed in blue shades, the matrix arm is displayed in green shades and the specific γ-carbonic anhydrase module is displayed in orange shades. Individual N, Q, P_D_ and P_P_ modules are indicated. **b** The resulting atomic model, each individual 45 proteins are annotated and displayed with a different color.

## Carbonic anhydrase: specific component of the plant complex I

The main feature of the plant complex I is the presence of an additional globular matrix-exposed domain, bound to the membrane arm. It holds an overlapping position with the NDUFA10/42-kDa subunit found in mammalian complex I^4^. This domain is formed by a trimeric complex, the carbonic anhydrase (CA). CA are zinc-containing enzymes that catalyze the reversible hydration of CO_2_ to HCO_3_^-^, whose roles are usually carbon fixation and pH maintenance^17^. The carbonic anhydrase activity evolved independently several times during evolution, resulting in several CA enzyme families. The plant mitochondrial CA is part of the gamma-carbonic anhydrase (γCA) family, which was originally identified in the archaeon *Methanosarcina thermophila*^18^. Here, the overall structure of the γCA is conserved: it is a trimer composed of characteristic left-handed beta-helix monomers, forming triangular prisms (Fig. 2a). However, in contrast to the archaeal enzyme, the γCA present in plant complex I is an heterotrimer. Indeed, in Arabidopsis five γCA genes were identified. Three have highly conserved catalytic domains and are termed γCA1-3, similar to the archaeal enzyme, while the two other show less homology to their prokaryotic homologues especially in the catalytic domain and were thus termed γCA-like proteins^19^. Our reconstruction shows that the γCA domain is formed by one copy of γCAL1, representing the most proximal subunit of the membrane part, and two copies of γCA1 (Fig. 2a and Extended Data Fig. 5). Each subunit have long N and C-terminal extensions interconnecting the 3 subunits and contacting the membrane subunits of complex I (Fig. 2c). These extensions also serve as anchors for the γCA domain by interacting with the plant specific P2 protein N-terminal part which is anchored in the membrane. The C-terminal part of the canonical Nad6 protein also contacts the CA domain stabilizing it. γCA is an enzyme, where one histidine of one subunit and two histidine of the adjacent subunit contribute to coordinate a zinc atom. However, as γCAL proteins lack two of those three histidines, it impairs the formation of the zinc coordination with both γCA subunits. Hence the only conserved active site is at the interface of the two γCAs, which we confirmed by visualizing both the zinc and an additional density most likely corresponding to HCO_3_ ^-^ (Fig. 2b and Extended Data Fig. 5). This confirms that the γCA domain associated with complex I is indeed active. However, the function of complex I-associated γCA is not well understood. It was suggested to play a role in complex I assembly as γCA is found in early assembly intermediates of the complex and is essential for complex I formation^20^. Indeed, embryo development of γCA mutants is strongly delayed and seed development arrested before maturation. Moreover complemented mutants with inactive γCA variants is sufficient to complement the mutant phenotype, showing that carbonic anhydrase activity is not essential^19^. Recently it was shown that carbonic anhydrase activity is also present in photosynthetic complex I, but is carried out by a different type of CA enzyme, positioned at the end of the P_D_ module that would contribute to proton generation^21^. However, in our complex, the only conserved active site is positioned at the most distal position relative to the membrane. Nevertheless, one can’t exclude its contribution to proton generation, although its effect would be relatively modest.

**Figure 2.**
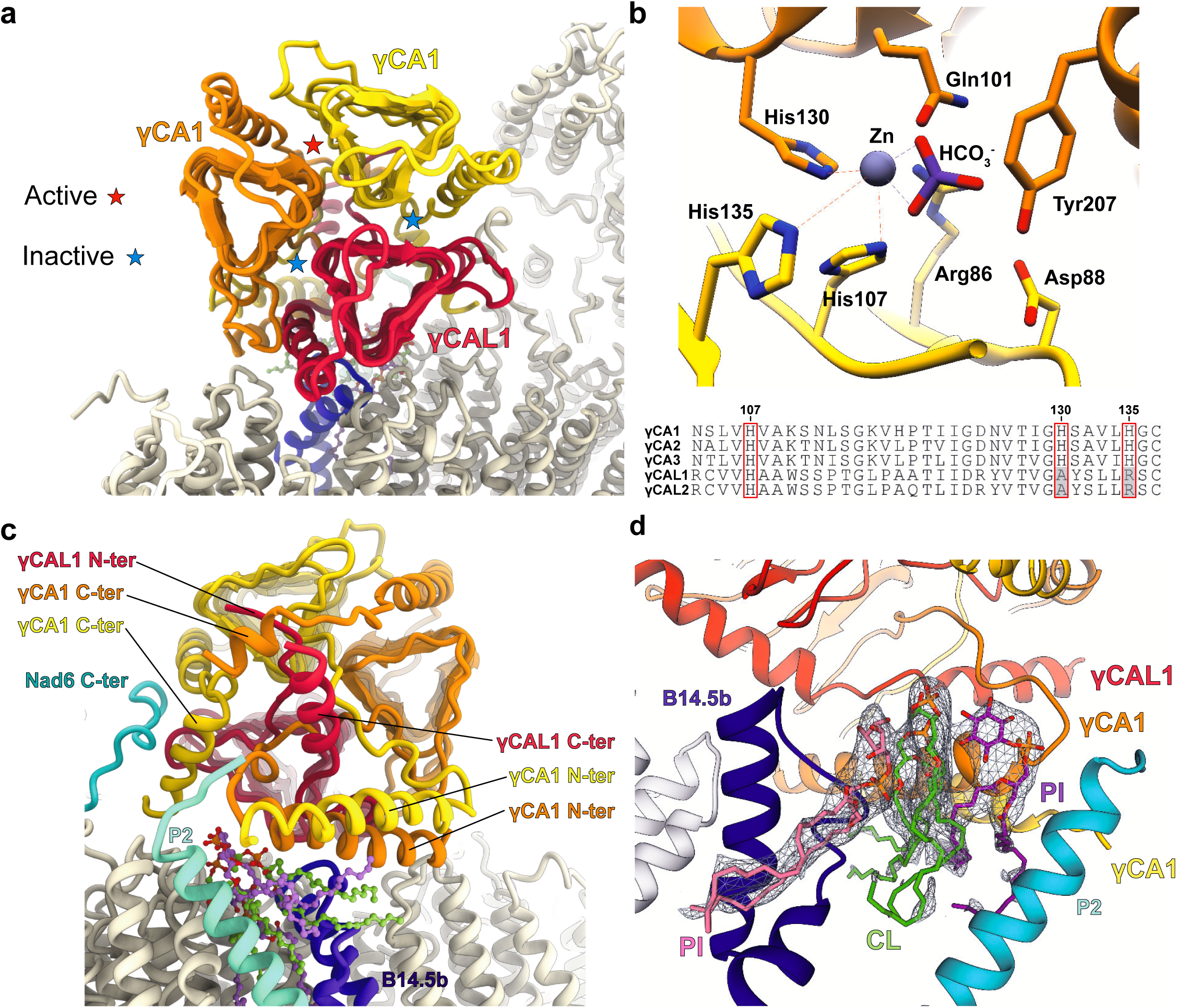
Hetero-trimeric γ-carbonic anhydrase is a specific component of the plant mitochondrial complex I. **a-c** Front and back view of the carbonic anhydrase showing individual subunits and contacting proteins. Active and inactive sites are shown in **a. b** Close up of the γCA1/γCA1 only conserved active site, showing the coordinate zinc by the three histidine residues between the two γCA1 subunits. Sequence alignment of γCA and γCAL highlight the catalytic histidines. **c** Back view of the carbonic anhydrase, highlighting the N and C-ter extensions of the γCA and γCAL as well as the proteins allowing to anchor the whole domain to the membrane arm. **d** The lipid block composed of one central cardiolipin and two phosphatidylinositol molecules is displayed in its density. The lipids are sandwiched between the carbonic anhydrase domain, the plant-specific protein P2, Nad2 and B14.5. Both inositol head groups contribute to coordinate the cardiolipin. CL stands for cardiolipin and PI for phosphatidylinositol.

Interestingly, the CA domain is in tight interaction with a peculiar lipid complex positioned below the latter and between the plant specific P2 and proteins B14.5b and Nad2, close to the Nad2-Nad4 junction (Fig. 2d). This lipid complex is composed of one cardiolipin coordinated by two phosphatidylinositol. Each inositol heads sandwich the glycerol linker of the cardiolipin and stabilize it. This lipid block was also observed in the assembly intermediate (see below) and appears to be very stable. Given the essential role of lipids for complex I activity, this lipid block might be crucial for the plant complex I efficiency and stability, which might be why CA mutants are so heavily affected^3,22^. Altogether, our analysis strongly suggest that the γCA is essential for the membrane part formation and stability, acting as an essential architectural factor that would remain even in mature complex.

## Assembly intermediate of the plant complex I

During 3D classification, two classes of complex I lacking the most distal proton pumping components of the membrane domain – Nad4 and Nad5 and their associated additional protein subunits – were found to naturally accumulate in our sample (Extended Data Fig. 1). One of the two classes presented an additional globular density on the intermembrane space exposed side of complex I. Such density wasn’t observed in full complex I classes. This class was thus attributed to an assembly intermediate of complex I (Fig. 3). In plants, such complex was already described to accumulate naturally. Indeed, extensive biochemical analyses identified complex I* as an assembly intermediate lacking the most distal membrane components and presenting an additional factor, GLDH (L-Galactono-1,4-lactone dehydrogenase). GLDH was identified in our mass spectrometry data (Extended Data Table 1), along with the other components of complex I. Likewise γCA, GLDH is an enzyme catalysing the last enzymatic step of the ascorbate biosynthetic pathway in plants^23^. It was never found in the full complex I, however, it was shown to associate with complex I intermediates, complex I* being the largest, as well as with smaller complex intermediates that were not observed here^20^. Moreover, in *gldh* knock-out mutant, complex I is undetectable, which hinted its role as a plant specific assembly factor^14,24^. Using focused classification and refinement, we were able to derive an intermediate-resolution reconstruction on the additional density corresponding to GLDH which was sufficient to attribute the density to the protein. Importantly, the complex I binding domain of GLDH is resolved at near-atomic resolution, thus validating our assignment (Fig. 3b-d). In this assembly intermediate, GLDH interacts with B14.5b - that is slightly displaced compared to the mature complex - and 15kDa. Moreover the whole carbonic anhydrase is slightly shifted toward the missing P_D_ module (Fig. 3c). We also successfully identified the N-terminal tail of GLDH bound to the maturing complex I, between B14.5b and 15kDa preventing the plant specific protein P1 from docking to the P_P_ module. P1 appears to functionally replace the NDUFB5 subunit, which is absent in plants. In the GLDH context the P1 path is blocked in the assembly intermediate due to B14.5b movement and N-ter tail of GLDH (Fig. 3). Thus, it appears that GLDH prevents the binding of P1 to the P_P_ module, until its release, which will then allow association with the P_D_ module.

**Figure 3.**
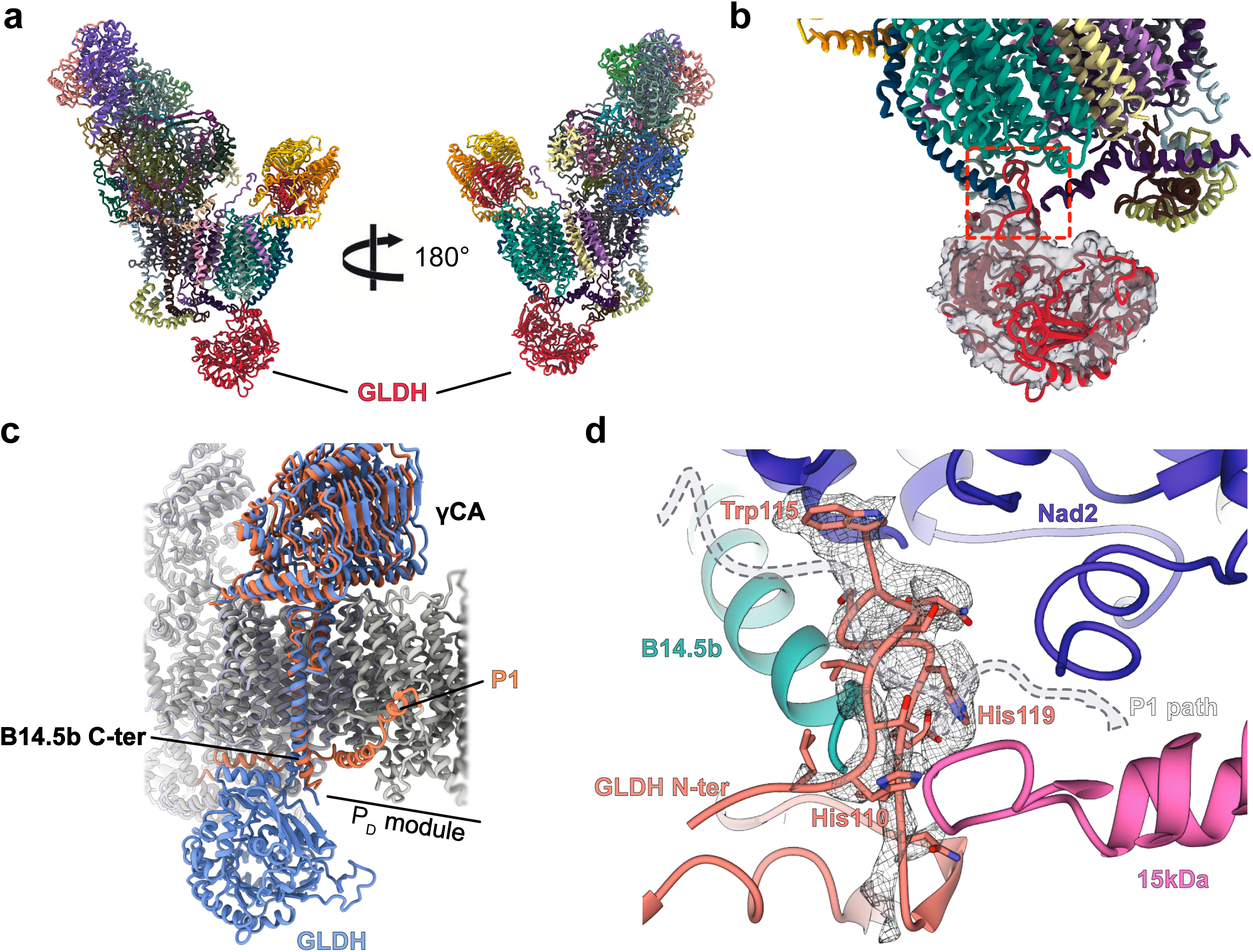
GLDH is a plant-specific assembly factor of mitochondrial complex I. **a** Atomic model of the assembly intermediate, the whole P_D_ module is missing and GLDH contacts the P_P_ module on the intermembrane side. **b** GLDH is shown in cut-out and filtered density. The interaction point is highlighted and a zoomed view is shown in **d. c** Overall movement in the assembly intermediate compared to the mature complex, B14.5b C-terminal part is slightly deviated and the whole γCA domain is shifted, GLDH and absence of the P1 protein are also highlighted. Coral correspond to mature complex I and blue to assembly intermediate. **d** Focused view of the interaction point of GLDH with the complex. The N-terminal part of GLDH was identified thanks to specific aromatic residues and is shown in its density. The N-ter part of GLDH as well as the deviation of B14.5b block P1 path which is shown in dashed lines.

In conclusion, our high-resolution cryo-EM structures of the plant mitochondrial complex I, reveal a most likely ancestral features of this respiratory complex (Fig. 4). Indeed, γCA was identified as a complex I component in all Viridiplantae, including algae of the Chlorophyceae class^11,12,25–27^, as well as in *Euglena gracilis*, a photosynthetic protozoan related to trypanosomes^28^ and possibly *Acanthamoeba castellanii*, a protozoan amoeboid^29^ and in non-photosynthetic organisms (e.g. Tetrahymena and Reclinomonas)^29^ (Fig. 4). Given the wide presence of the γCA in eukaryotes – members of the γCA are found in all eukaryotic lineages but not in opisthokonts – we suggest that γCA proteins may be an ancestral feature of mitochondrial complex I. Moreover we describe a yet unobserved plant-specific assembly intermediate, shedding light into the respiratory complex I maturation.

**Figure 4.**
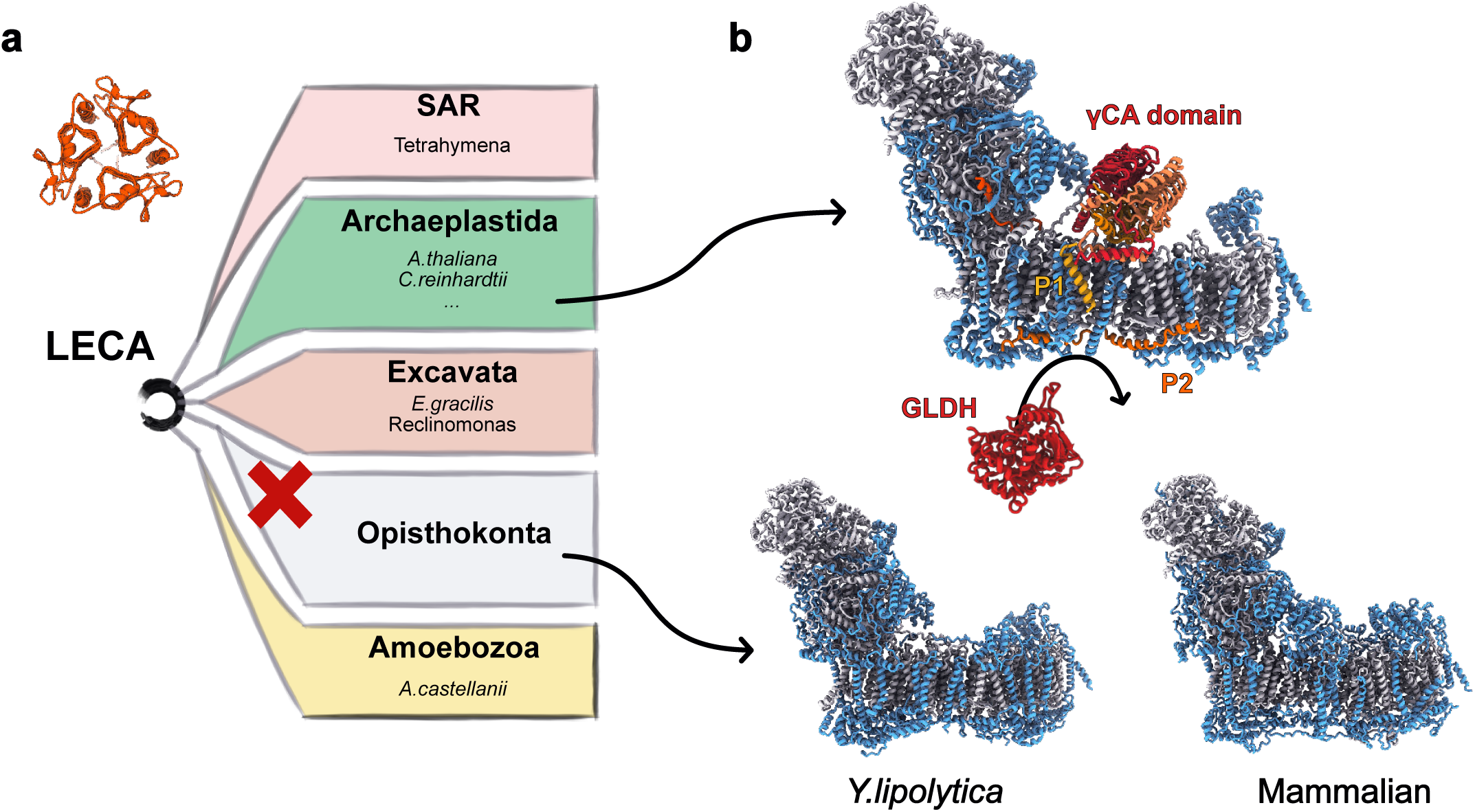
Plant mitochondrial complex I specific subunits. **a** Schematic representation of the five major eukaryote lineages. Organisms where γ-carbonic anhydrase is hypothesized to be part of mitochondrial complex I are listed next to their respective lineage. The red cross indicate the loss of mitochondrial γ-carbonic anhydrase in opisthokonta. **b** Comparison of mitochondrial complex I structures. Core subunits are shown in light gray and supernumary subunits in light blue. Plant specific proteins are highlighted in red-orange colours. *Y.lipolytica* (PDB:6RFR) and mammalian (PDB:6G2J) complex I don’t have the γ-carbonic anhydrase domain.

## METHODS

Methods and any associated references are available in the online version of the paper.

## ACKNOWLEDGEMENTS

This work has benefitted from the facilities and expertise of the Biophysical and Structural Chemistry platform (BPCS) at IECB, CNRS UMS3033, Inserm US001, University of Bordeaux. We thank A. Bezault for assistance with the Talos Arctica electron microscope. We thank, J. Chicher and P. Hamman of the Strasbourg Espanade proteomic analysis for the proteomic analysis. This work was supported by a European Research Council Starting Grant (TransTryp ID:759120) and a Agence Nationale de la Recherche (ANR) grant [MITRA, ANR-16-CE11-0024-02]] to YH.

## AUTHOR CONTRIBUTIONS

FW, HS and YH designed and coordinated the experiments. FW purified the mitochondria and mitochondrial complex I. HS and CP prepared grids. HS acquired the cryo-EM data and processed the cryo-EM results with FW. HS and FW built the atomic models. LK performed the mass-spectrometry experiments. FW, HS and YH interpreted the structure. FW, HS, CP and YH wrote and edited the manuscript.

## DATA AVAILABILITY

The cryo-EM maps of plant mitochondrial complex I have been deposited at the Electron Microscopy Data Bank (EMDB): EMDB-XXXXX. The corresponding atomic models been deposited in the Protein Data Bank (PDB) under the accession XXXX. Mass spectrometric data were deposited with the ProteomeXchange Consortium via the PRIDE partner repository with the dataset identifier XXXXXXX.

---

**Extended Data Figure 1.**
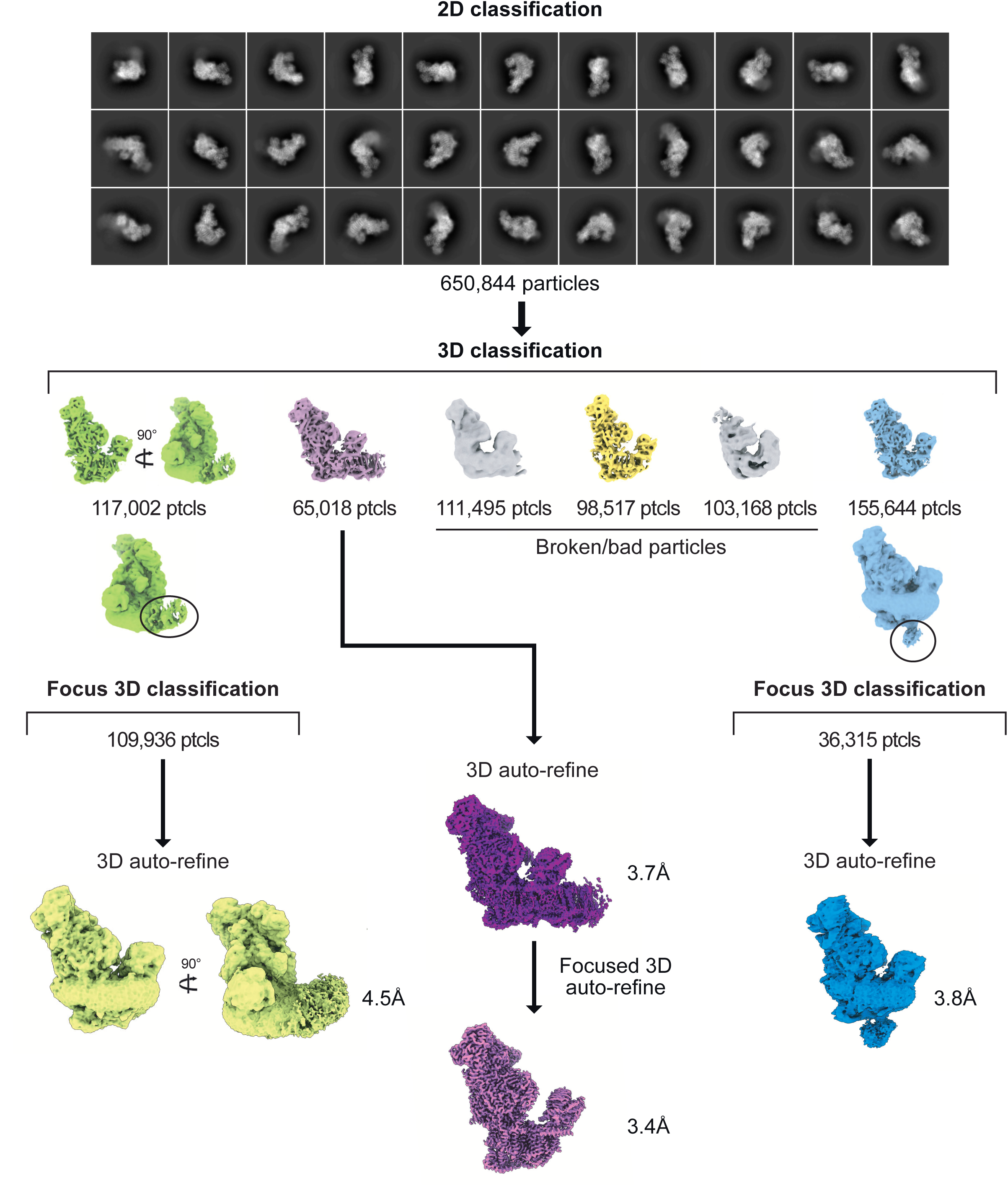
Data processing workflow. Graphical summary of the processing workflow described in Methods, with 2D classes presented in **a** and 3D processing and refinement presented in **b**. Density in purple represent the mature and focused reconstructions, density in blue represent the assembly intermediate. The green density maps correspond to a complex where the whole P_D_ module is shifted, but that complex was not further analyzed.

**Extended Data Figure 2.**
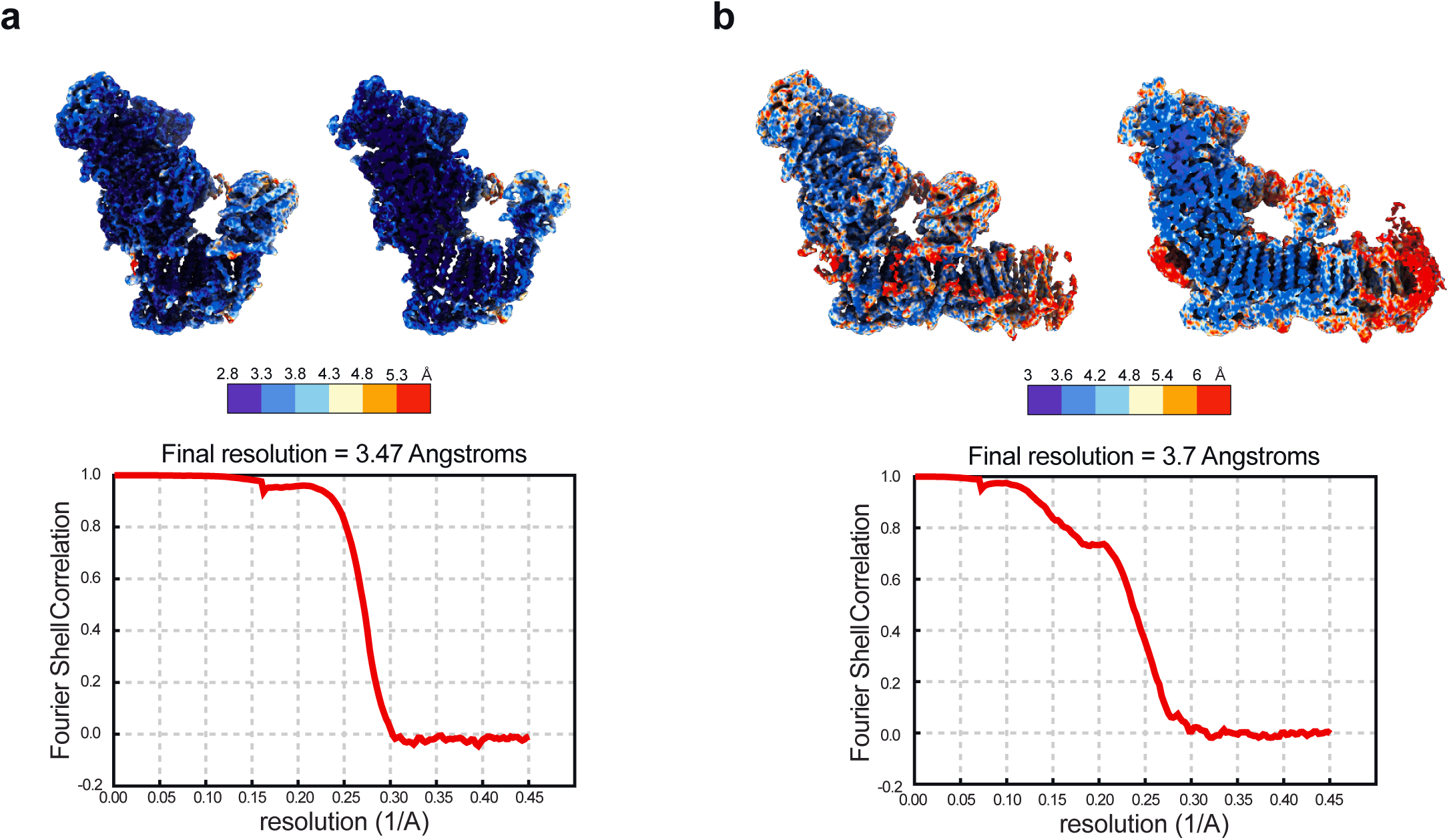
Local resolution of the final reconstructions. Local resolutions of both the full **a** and focused **b** complex I are shown. The maps are colored by resolution, generated using ResMap^30^. Maps are also shown in cut view. For both reconstructions FSC plots are displayed for resolution estimation. The map resolution is calculated at the 0.143 threshold.

**Extended Data Figure 3.**
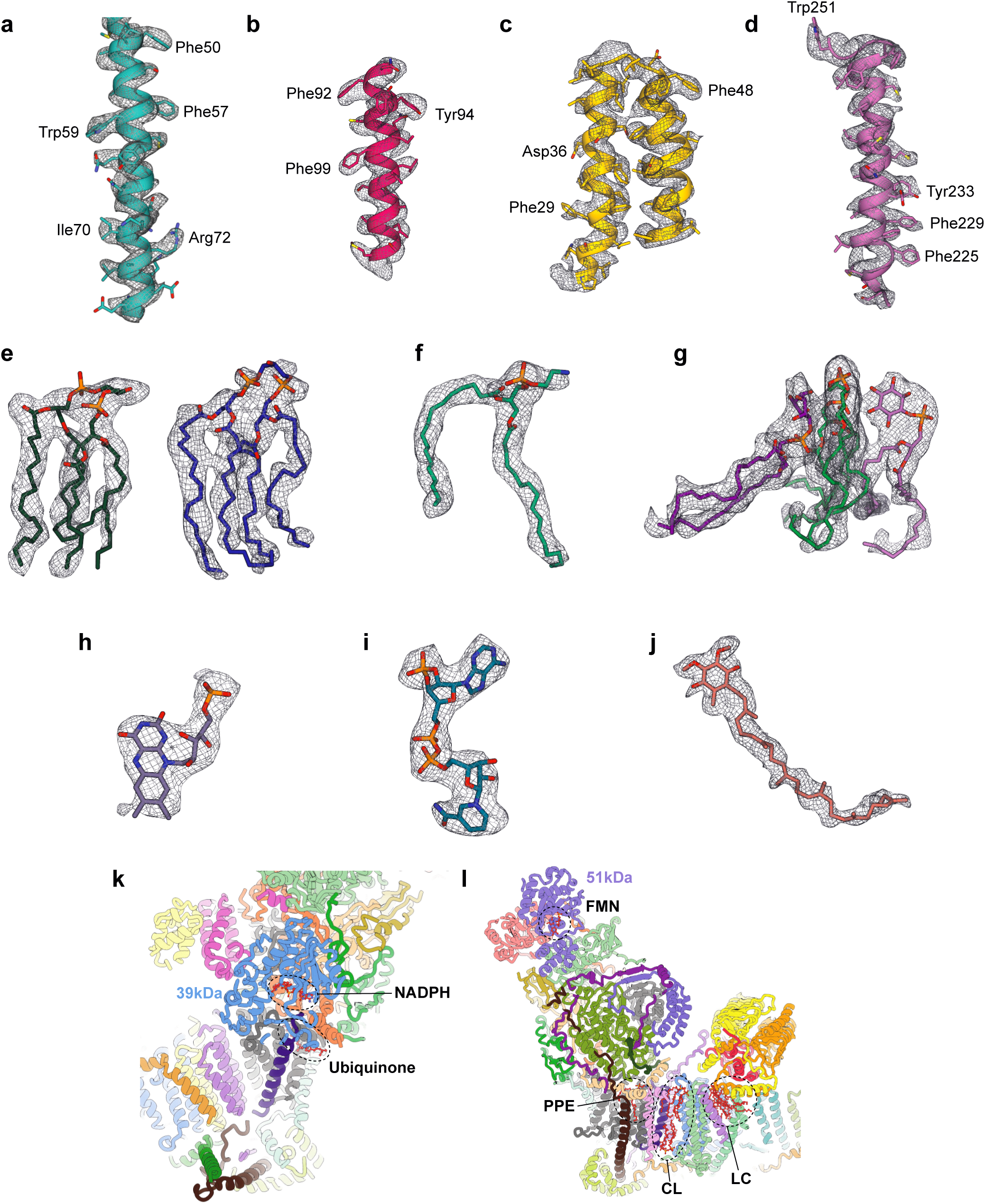
Representative cryo-EM densities. **a-d** Proteins densities with **a** segment of protein B16.6, **b** segment of protein Nad2, **c** segment of protein Nad6 and **d** segment of protein Nad1. **e-f** Lipids identified in their respective densities with **e** cardiolipins, **f** phosphatidylethanolamine and **g** the carbonic anhydrase lipid complex with the cardiolipin in green and phosphatidylinositols in purple and orchid. **h-j** Additionnal ligands with **h** FMN of 51kDa subunit, **i** NADPH of the 39kDa subunit and **j** the ubiquinone. Positions of the ligands are indicated on the atomic model on **k** and **l** with PPE for phosphatidylethanolamine, CL for cardiolipin and LC for lipid complex.

**Extended Data Fig4.**
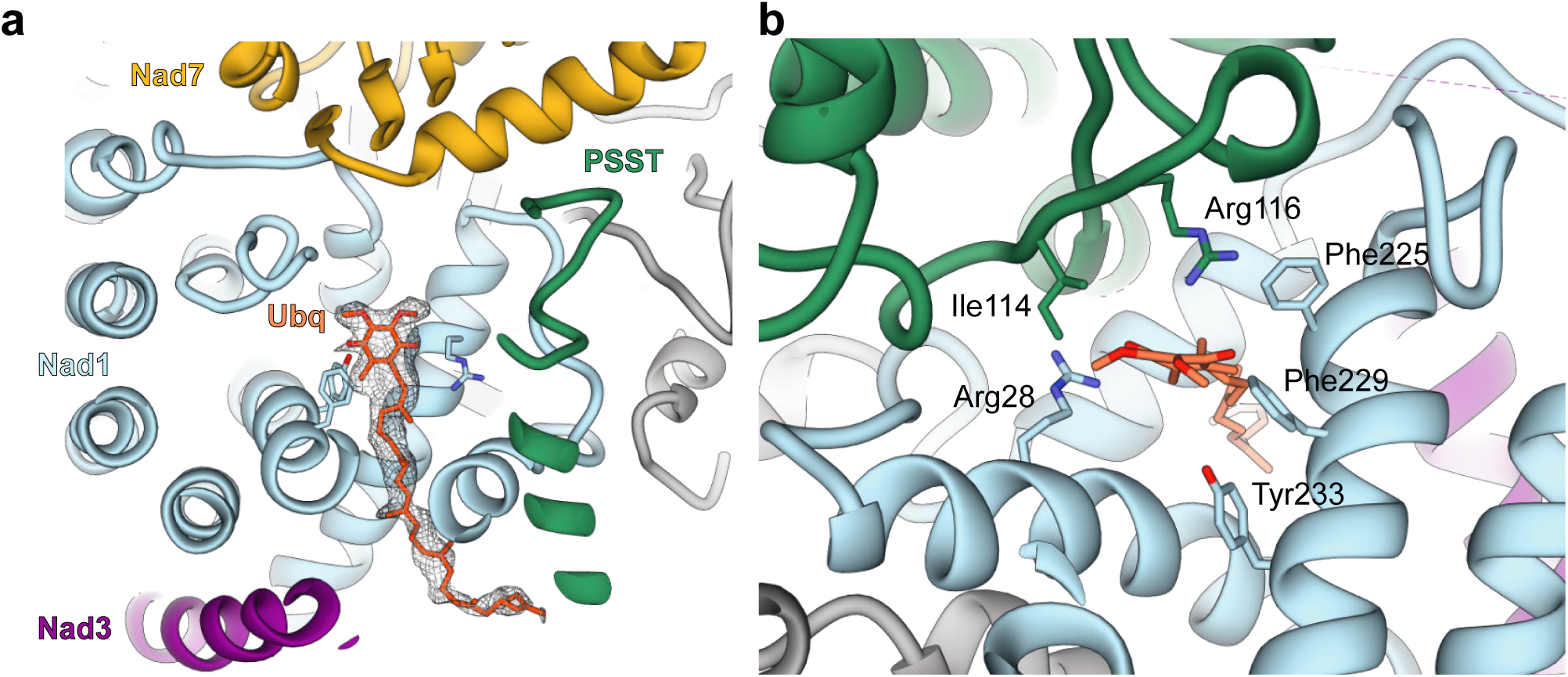
Ubiquinone binding site. **a** Overall view of the ubiquinone pocket mainly formed by Nad1 and PPST. The ubiquinone is shown in its density. **b** Focus on the ubiquinone head group and surrounding residues. The position and coordination of ubiquinone are similar to what has been observed in the *Y.lipolytica* mitochondrial complex I^3^.

**Extended Data Figure 5.**
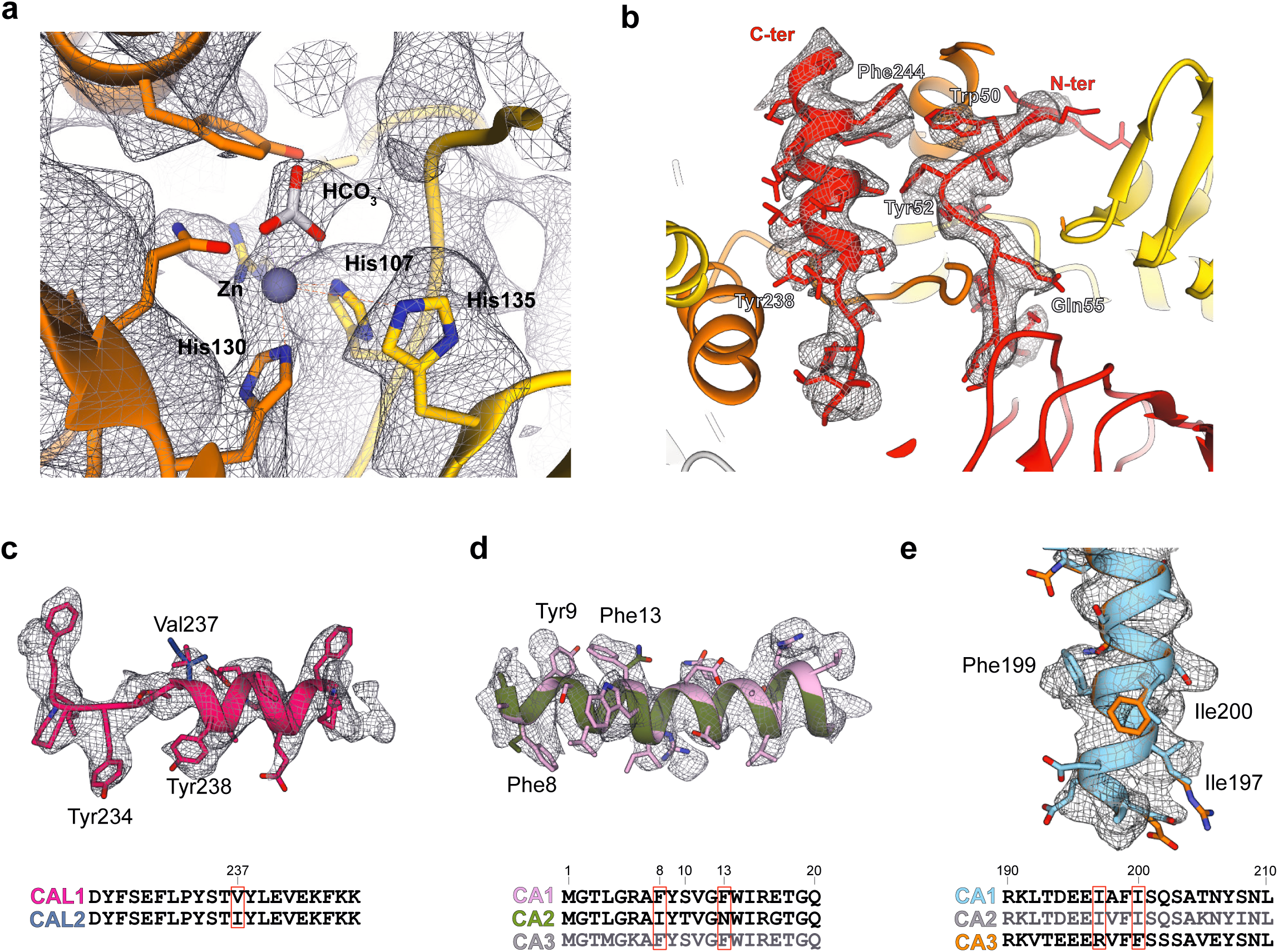
Identification of carbonic anhydrase subunits. **a** Density of the zinc and the HCO_3_^-^ are shown, as well as the surrounding residues. The distinction between γCA and γCAL was done based on N and C-ter tails as well as conserved histidine residues, identification of the γCAL is shown in **b**. Models in their respective densities and sequence alignment are presented for the part that contributed to identify each of the subunit. Both possibilities are displayed and alignments are color-coded in accordance to the models. **c** To differentiate between γCAL1 and γCAL2 Val237 was used, which would be an isoleucine in γCAL2. However, given the high similarity of the two proteins (only 5 different aa excluding the mitochondrial target sequence) and previous genetic screens^19^ both could be there. **d** For the γCA subunits, for chain p, residues Phe8 and Phe13 allowed to determine that it was γCA1 or 3 and not γCA2. **e** Residue Ile200 and Ile197, which would be a Phe and an Arg in γCA3, were used to distinguish between γCA1 and γCA3, confirming that the subunit is indeed γCA1. Still for the second γCA (chain q) the scant density did not allow to clearly distinguish between γCA1 and γCA2, and was therefore built as γCA1.

**Extended Data Figure 6.**
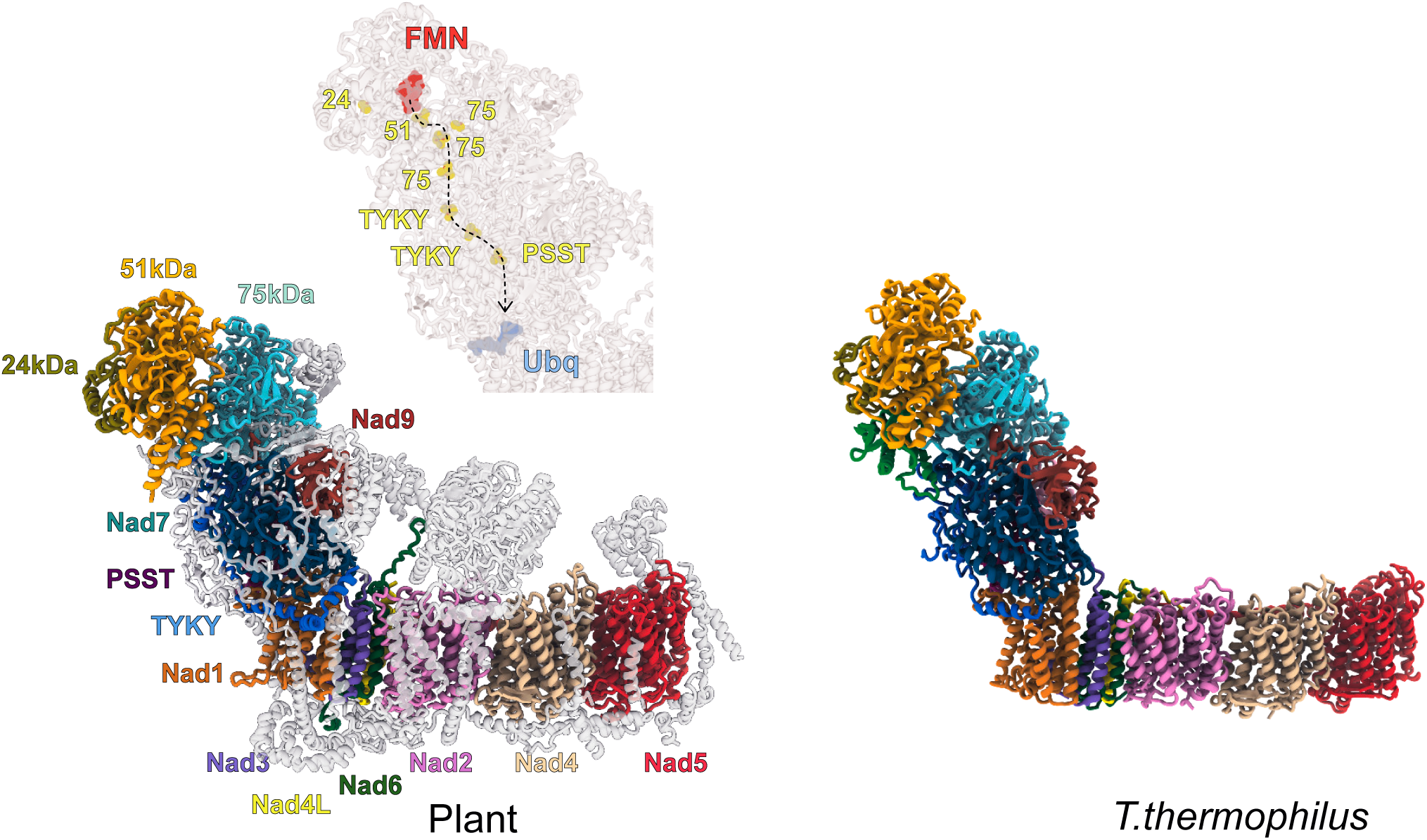
Plant mitochondrial complex I comparison with prokaryotic complex I. Plant mitochondrial complex I is shown compared to its bacterial homologue (*T.thermophilus* model PDB:4HEA). The 14 core subunits are colored in different shades and their names are indicated. Mitochondria and plant specific proteins are shown in gray and transparent. Electron path in matrix arm from FMN to ubiquinone is also shown for the plant complex I.

## MATERIAL & METHODS

### Mitochondrial Complex I purification

Cauliflower (*Brassica oleracea* var. botrytis) mitochondria were purified as previously described^13^. Quickly, fresh cauliflower inflorescence tissue was blended in extraction buffer containing 0.3 M mannitol, 30 mM sodium pyrophosphate (10.H_2_O), 0.5 % BSA, 0.8 % (w/v) polyvinylpyrrolidone-25, 2mM beta-mercaptoethanol, 1 mM EDTA, 20 mM ascorbate and 5 mM cysteine, pH 7.5. Lysate was filtered and clarified by centrifugation at 1.500 g, 10 min at 4°C. Supernatant was kept and centrifuged at 18.000 g, 15 min at 4°C. Organelle pellet was re-suspended in wash buffer (0.3 M mannitol and 10 mM phosphate buffer, 1 mM EDTA, pH 7.5) and the precedent centrifugations were repeated once. The resulting organelle pellet was re-suspended in wash buffer and loaded on a single-step 30% Percoll gradient (in wash buffer without EDTA) and run for 1h30 at 40.000 g. Mitochondria were retrieved, washed two times and flash frozen in liquid nitrogen.

For complex I purification, mitochondria were re-suspended in Lysis buffer (20 mM HEPES-KOH, pH 7.6, 100 mM KCl, 1 mM DTT, 2% n-DDM, 1mM EDTA, supplemented with proteases inhibitors (Complete)) incubated for 15 min in 4°C. Lysate was clarified by centrifugation at 30.000 g, 20 min at 4°C. The supernatant was loaded on a 10-50% sucrose gradient buffer (20 mM HEPES-KOH, pH 7.6, 50 mM KCl, 1 mM DTT, 0.2% n-DDM, 1mM EDTA, supplemented with proteases inhibitors (Complete)) and run for 16 h at 75,000 g. The fraction corresponding to complex I was collected, pelleted and re-suspended in final resuspension buffer (same as sucrose gradient buffer without sucrose and only 0.1% n-DDM).

### Grid preparation

4 µL of the samples at a concentration of 1 µg/µl was applied onto Quantifoil R2/2 300-mesh holey carbon grid, coated with thin home-made continuous carbon film and glow-discharged. The sample was incubated on the grid for 30 sec and then blotted with filter paper for 2.5 sec in a temperature and humidity controlled Vitrobot Mark IV (T = 4°C, humidity 100%, blot force 5) followed by vitrification in liquid ethane.

### Single particle cryo-electron microscopy data collection

Data collection was performed on a Talos Arctica instrument (Thermofisher Company) at 200 kV using the SerialEM software for automated data acquisition. Data were collected at a nominal underfocus of - 0.5 to −2.5 µm at a magnification of 36,000 X yielding a pixel size of 1.2 Å. Micrographs were recorded as movie stack on a K2 direct electron detector (GATAN Company), each movie stack were fractionated into 65 frames for a total exposure of 1 sec corresponding to an electron dose of 45 ē/Å2.

### Electron microscopy image processing

Drift and gain correction and dose weighting were performed using MotionCorr2^31^. A dose weighted average image of the whole stack was used to determine the contrast transfer function with the software Gctf^32^. The following process has been achieved using RELION 3.0^15^. Particles were picked using the general model of crYOLO 1.5 ^33^. Output box files from crYOLO were imported in RELION and particles were extracted. After 2D classification, 650,844 particles were extracted with a box size of 360 pixels and binned threefold, resulting in a 120 pixels bow size four fold for 3D classification into 6 classes for each subunits (Extended Data Fig. 1). One subclass depicting high-resolution features have been selected for the mature complex I refinement with 65,018 particles. After Bayesian polishing a focused refinement has been performed using mask excluding the P_D_ module yielding respectively 3.7 and 3.47Å resolution. A second subclass containing the maturation factor GLDH composed of 155,644 particles was selected for further focused 3D classification on the GLDH area. The 36,513 particles of GLDH enriched subclass were extracted for refinement and reached 3.8Å. Determination of the local resolution of the final density map was performed using ResMap^30^.

### Structure building and model refinement

The atomic model of the plant mitochondrial complex I was built into the high-resolution maps using Coot, Phenix and Chimera. Atomic models from *Y.lipolytica* (6RFR) and *M.musculus* (6G2J) were used as starting points for protein identification and modelisation. Similarly to Waltz & Soufari 2019^13^, as the genome of cauliflower is not sequenced, and the closest fully sequenced member of the family (*Brassica oleracea* subsp. oleracea) is poorly annotated, to facilitate comprehension and analysis we positioned Arabidopsis proteins in the cauliflower map. The online SWISS-MODEL service was used to generate initial models for bacterial and mitochondria conserved r-proteins. Models were then rigid body fitted to the density in Chimera^34^ and all subsequent modeling was done in Coot^35^. The global atomic model was refined with VMD using the Molecular Dynamic Flexible Fitting (MDFF) then with PHENIX using a combination of real and reciprocal space refinement for proteins.

### Proteomic and statistical analyses of mitochondrial ribosome composition

Mass spectrometry analyses of the ribosome fractions were performed at the Strasbourg-Esplanade proteomic platform and performed as previously^36^. In brief, proteins were trypsin digested, mass spectrometry analyses and quantitative proteomics were carried out by nano LC-ESI-MS/MS analysis on AB Sciex TripleTOF mass spectrometers and quantitative label-free analysis was performed through in-house bioinformatics pipelines. Data were searched against the TAIR *A. thaliana* database with a decoy strategy (release TAIRv10, 27281 forward protein sequences), and home-made cauliflower database extracted from UniProtKB (Swissprot+TrEMBL) including Brassica sub-taxonomy.

### Figure preparation

Figures featuring cryo-EM densities as well as atomic models were visualized with UCSF ChimeraX^37^ and Chimera^34^.

